# Efficient reconciliation of genomic datasets of high similarity

**DOI:** 10.1101/2022.06.07.495186

**Authors:** Yoshihiro Shibuya, Djamal Belazzougui, Gregory Kucherov

## Abstract

We apply Invertible Bloom Lookup Tables (IBLTs) to the comparison of *k*-mer sets originated from large DNA sequence datasets. We show that for similar datasets, IBLTs provide a more space-efficient and, at the same time, more accurate method for estimating Jaccard similarity of underlying *k*-mer sets, compared to MinHash which is a go-to sketching technique for efficient pairwise similarity estimation. This is achieved by combining IBLTs with *k*-mer sampling based on syncmers, which constitute a context-independent alternative to minimizers and provide an unbiased estimator of Jaccard similarity. A key property of our method is that involved data structures require space proportional to the difference of *k*-mer sets and are independent of the size of sets themselves. As another application, we show how our ideas can be applied in order to efficiently compute (an approximation of) *k*-mers that differ between two datasets, still using space only proportional to their number. We experimentally illustrate our results on both simulated and real data (*SARS-CoV-2* and *Streptococcus Pneumoniae* genomes).

Available at: https://github.com/yhhshb/km-peeler.git

## 1 Introduction

Alignment-free methods became a prevalent paradigm in computational analysis of modern genomic datasets. However, despite being faster than their alignment-based counterparts, algorithms based on *k*-mer sets are starting to struggle when applied to the large datasets produced nowadays [20, 13, 16]. To deal with this issue, a considerable effort has been put to developing optimized data structures, with succinct solutions [25, 23, 16] and approximate membership data structures [29, 13, 2, 14, 3] being two examples.

In recent years, sketching techniques have been gaining increasing attention thanks to their capacity of drastically decreasing space usage. MinHash is probably the most well-known representative of this family of algorithms. Application of MinHash to comparison of DNA sequence datasets was pioneered in Mash software [24] and subsequently used in several other tools. With this approach, input datasets are transformed into smaller “sketches” on which subsequent comparisons are performed. In short, sequences are first fragmented into their constituent *k*-mers which are then hashed, with each sketch storing only *s* minimum values, with *s* defined by the user. The fraction of shared hashes between two sketches is an unbiased estimator of the Jaccard similarity index [4]. A MinHash sketch can thus be viewed as a sample of the set of *k*-mers of the sequence it represents. Given that *s* is much smaller than the genome length, working with the sampled hashes leads to fast pairwise comparisons using small memory. However, when two sequences are close and share most of their *k*-mers, MinHash sketches of small size are not able to reliably estimate their degree of similarity since differences are likely to be missed during sampling.

In this work, we propose an alternative approach to evaluate the difference in *k*-mer composition of two related datasets. Our method relies on Invertible Bloom Lookup Table (IBLT) data structure [12, 10] which is an extension of Bloom filters, supporting deletions of items and, most importantly, enumeration (with high probability) of stored items. One of the applications of IBLT is reconciliation of two sets of items: in a scenario considered in [12], a set *A* is stored in an IBLT which is then transmitted to the holder of another set *B*. By screening *B* against the IBLT of *A* it is possible to recover the items *A* \ *B* and *B* \ *A*, with high probability. This is done through the so-called *peeling* procedure [7].

In this paper we make one step further: inspired by ideas of [26], we recover both *A* \ *B* and *B* \ *A* from IBLTs of *A* and *B*, rather than from an IBLT of one of them and the whole other set. Furthermore, a crucial property is that the *size of these IBLTs is bounded in terms of the symmetric difference size* (*A* \ *B*) ∪ (*B* \ *A*) rather than the size of the original sets. This provides a key to the efficiency of our solution when input sets are similar: even if input sets are very big, their difference can be recovered using a data structure (sketch) whose size is proportional to the size of the difference of those sets rather than of the sets themselves. Estimating the symmetric difference allows us to estimate the Jaccard similarity, using information about the sizes of input sets. Thus, whereas close datasets require larger MinHash sketches to be properly compared, our method, on the contrary, requires smaller memory.

Another ingredient of our solution is *k*-mer sampling. Intuitively, since two adjacent *k*-mers share *k* − 1 bases, the information stored in the set of all *k*-mers appears highly redundant. One popular method of sampling *k*-mers from genomic sequences is based on minimizers [28]. Under this technique, consecutive sampled *k*-mers are within a bounded distance from each other and therefore no large portion of the sequence can remain unsampled. Another favorable property is that similar regions are likely to yield similar samples of minimizers. However, it has recently been shown that estimating Jaccard similarity based on minimizer sampling leads to a bias [1]. Here we propose to replace minimizers by syncmers [8]. Syncmers provide another way of *k*-mer sampling which has certain advantages over minimizers. As opposed to minimizers, syncmers are not context-dependent: for a *k*-mer to be a syncmer depends on the *k*-mer alone regardless the context where it occurs, and, under standard randomness assumptions on involved hash functions, all *k*-mers have equal chance to be a syncmer. As a consequence, syncmer sampling leads to an unbiased estimate of Jaccard similarity, as the fraction of syncmers among shared *k*-mers (intersection) is expected to be the same as that among all *k*-mers (union). We experimentally validate that this is indeed the case.

By combining syncmer sampling with IBLTs, we obtain a space-efficient method for accurately estimating Jaccard similarity for similar datasets. For datasets of high similarity, the proposed method is superior to the popular MinHash algorithm [24], both in terms of memory and precision. We also propose an application of this technique to retrieve *k*-mers that differ between two given datasets. Our method computes a superset of those *k*-mers with a limited number of spurious *k*-mers. In particular, under the assumption that each *k*-mer occurs once, our method computes the exact set differences between involved *k*-mer sets. We validate our algorithms on both simulated data and on real datasets made of *SARS-CoV-2* and *Straphilococcus Pneumoniae* genomes.

## 2 Technical preliminaries

We consider DNA alphabet Σ = {*A, C, G, T*} even though our algorithms can be easily generalized. Given a string *S* ∈ Σ^*∗*^, we use the notation *S*[*i, k*] to indicate the substring of length *k* starting at position *i* called a *k*-mer. The *k*-mer set *K*_*S*_ of *S* is the set of *k*-mers *S*[*i, k*] for *i* ∈ [0, |*S*| − *k* + 1].

### 2.1 Minimizers

Independently introduced in [28] and [30], minimizers are defined by a triplet of parameters (*k, w, h*), where *k* is the *k*-mer length, *w* a window size, and *h* a function defining an order on *k*-mers. *h* is usually chosen to be an appropriately defined hash function, the lexicographical order is rarely used in practice due to its poor statistical properties.

Each window *S*[*i, w* + *k* − 1] defines a minimizer which is the minimal *k*-mer among *w k*-mers occurring in *S*[*i, w* + *k* − 1] w.r.t. the order given by *h*. Two neighboring minimizers are thus separated by at most *w* positions making it impossible to have large stretches of the original sequence not covered by any minimizers.

Since two neighboring windows at positions *i* and *i* + 1 are likely to share their minimizer, minimizers provide a way to sample *k*-mers from a sequence with bounded distance between consecutive sampled *k*-mers. An advantage of this sampling strategy is that similar sequences will likely have similar lists of minimizers, which makes it useful for mapping algorithms [19, 15]. Under reasonable assumptions, the density of minimizers, i.e. fraction of sampled *k*-mers, is 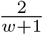 [28, 8]. If minimizer positions in the original sequence are not important, they can be discarded and the resulting *k*-mer multiset can be reduced to a simple *k*-mer set.

### 2.2 Syncmers

Minimizers are susceptible to mutations of any base of their window [8]. That is, a *k*-mer may cease to be a minimizer if a modified base occurs not only inside this *k*-mer, but also in its close neighborhood. Sampling with a higher density alleviates this problem but it reduces the advantages of the methods because more minimizers are selected. Methods to generate minimizer indices with the best possible density exist [6, 9] but they are usually offline algorithms, limiting their potential applications outside alignment.

Syncmers are a family of alternative methods to minimizers that does not suffer from this issue [8]. Similarly to minimizers, syncmers are defined using a triplet of parameters (*k, z, h*) where *z < k* is used to decompose each *k*-mer into its constituent *z*-mers and *h* defines an order over them. A *k*-mer *q* is a *syncmer* (called *closed syncmers* in [8]) iff its minimal *z*-mer occurs as a prefix (position *i* = 0) or as a suffix (position *i* = *k* − *z* + 1) of *q*. Thus, a syncmer is defined by its sequence alone, regardless the context in which it occurs. For this reason, syncmer sampling has been shown to be more resistant to mutations and then to improve the sensitivity of alignment algorithms [8].

Similar to minimizers, consecutive syncmers occur at a bounded distance. More precisely, consecutive syncmers must overlap by at least *z* characters and therefore “pave” the sequence without gaps. The fraction of syncmers among all *k*-mers is estimated to be 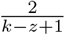 [8].

### 2.3 Invertible Bloom Lookup Tables

Invertible Bloom Lookup Tables (IBLT) [10, 12] are a generalization of Bloom filters for storing a set of elements (keys), drawn from a large universe, possibly associated with attribute values. In contrast to Bloom filters, in addition to insertions, IBLTs also support deletion of keys as well as listing. The latter operation succeeds with high probability (w.h.p.) depending on the number of stored keys relative to the size of the data structure. An important property is that this probability depends only on the number of keys stored at the moment of listing, and not across the entire lifespan of the data structure. Thus, at a given time, an IBLT can store a number of keys greatly exceeding the threshold for which it was built, returning to be fully functional whenever a sufficient number of deletions has taken place. Note also that IBLTs, in their basic version, don’t support multiple insertions of the same key.

An IBLT is an array *T* of *m* buckets together with *r* hash functions *h*_1_, …, *h*_*r*_ mapping a key universe *U* (in our case, *k*-mers or strings) to [0..*m* − 1] and an additional global hash function *h*_*e*_ on *U*. Each bucket *T* [*i*], *i* ∈ [0..*m* − 1], contains three fields: a counter *T* [*i*].*C*, a key field *T* [*i*].*P* and a hash field *T* [*i*].*H*, where *C* counts the number of keys hashed to bucket *i, P* stores the XOR-sum of the keys (in binary representation) hashed to bucket *i*, and *H* contains the XOR-sum of hashes produced by *h*_*e*_ on keys.

Adding a key *p* to the IBLT is done as follows. For each *j* ∈ {1, …, *r*}, we perform *T* [*h*_*j*_(*p*)].*C* = *T* [*h*_*j*_(*p*)].*C* + 1, *T* [*h*_*j*_(*p*)].*P* = *T* [*h*_*j*_(*p*)].*P* ⊕ *p*, and *T* [*h*_*j*_(*p*)].*H* = *T* [*h*_*j*_(*p*)].*H* ⊕ *h*_*e*_(*p*), where ⊕ stands for XOR. Given that XOR is the inverse operation of itself, deletion of *p* is done similarly except that *T* [*h*_*j*_(*p*)].*C* = *T* [*h*_*j*_(*p*)].*C* − 1.

Listing the keys held in an IBLT is done through the process of peeling working recursively as follows. If for some *i* we have *T* [*i*].*C* = 1, payload field *T* [*i*].*P* is supposed to contain a single key *p*. Field *H* is not strictly necessary, it acts as a “checksum” to verify that *p* is indeed a valid key by checking if *h*_*e*_(*T* [*h*_*j*_(*p*)].*P*) = *T* [*h*_*j*_(*p*)].*H*. This check is used to avoid the case when *T* [*i*].*C* = 1 whereas *T* [*i*].*P* is not a valid key, which can result from extraneous deletions of keys not present in the data structure. In Section 3.2 we will elaborate on the role of this field in our framework. If the check holds, key *p* can be reported and deleted (peeled) from the IBLT. Updating hash sums and counters is done in a similar way: *T* [*h*_*j*_(*p*)].*H* = *T* [*h*_*j*_(*p*)].*H* ⊕ *h*_*e*_(*p*) and *T* [*h*_*j*_(*p*)].*C* = *T* [*h*_*j*_(*p*)].*C* − 1. The procedure continues until all counters *T* [*i*].*C* are equal to zero.

At each moment, an IBLT is associated to a *r*-hypergraph where nodes are buckets and edges correspond to stored keys with each edge including the buckets a key is hashed to. Listing the keys contained in an IBLT then relies on the peelability property of random hypergraphs [7, 22]. Assume our hash functions are fully random. Then it is known that for *r* ≥ 3, a random *r*-hypergraph with *m* nodes and *n* edges is peelable w.h.p. iff *m* ≥ *c*_*r*_*n* where *c*_*r*_ is a constant peelability threshold. The first values of *c*_*r*_ are *c*_3_ ≈ 1.222, *c*_4_ ≈ 1.295, *c*_5_ ≈ 1.425, · · · [12]. Thus, allocating

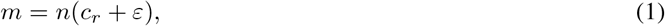

buckets, for *ε >* 0, for storing *n* keys guarantees successful peeling with high probability.

### 2.4 MinHash sketching

MinHash sketching was introduced in [4] as a method to estimate Jaccard similarity between two sets, applied to document comparison. In bioinformatics, MinHash was first applied in Mash software [24] and then successfully used in a number of other tools. Assume we are given a universe *U* and an order on *U* defined via a hash function *h*. For a set *A* ⊂ *U*, the bottom-*s* MinHash sketch of *A*, denoted 𝕊(*A*), is the set of *s* minimal elements of *A* (or their hashes), where *s* is a user-defined parameter. The Jaccard similarity index between two sets *A* and *B, J* (*A, B*) = |*A* ∩ *B*|*/*|*A* ∪ *B*|, can then be estimated from the sketches of *A* and *B*, namely

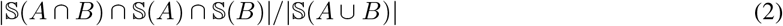

is an unbiased estimator of *J* (*A, B*).

The Jaccard similarity between the *k*-mer sets of two dataset constitutes a biologically relevant measure of their similarity. In particular, if involved datasets are genomic sequences, this measure allows one to estimate the mutation rate between the sequences [11, 24].

## 3 Methods

### 3.1 Set reconciliation from two IBLTs

Invertible Bloom Lookup Tables can be used to achieve set reconciliation between two sets *A* and *B*, that is to recover sets *A* \ *B* and *B* \ *A*. Under a scenario described in [12], the holder of *A* stores it in an IBLT *T*_*A*_ which is then transmitted to the holder of *B*. Elements of *B* are then deleted from *T*_*A*_. In the resulting IBLT, *P* -fields with *T*_*A*_[*i*].*C* = 1 correspond to elements of *A* \ *B* and those with *T*_*A*_[*i*].*C* = −1 to *B* \ *A*. The peeling process is applied to either of such fields. Whenever *T*_*A*_[*i*].*C* = 1, we delete *p* = *T*_*A*_[*i*].*P* from *T*_*A*_ on condition that *h*_*e*_(*p*) = *T*_*A*_[*i*].*H*. Similarly, whenever *T*_*A*_[*i*].*C* = −1, we add (XOR) *p* = *T*_*A*_[*i*].*P* to *T*_*A*_ on condition that *h*_*e*_(*p*) = *T*_*A*_[*i*].*H*. The process lists all elements of both *A* \ *B* and *B* \ *A* w.h.p.

Inspired by work [26], we modify the above scheme in order to recover the symmetric difference between *A* and *B* from their respective IBLTs *T*_*A*_ and *T*_*B*_, rather than from the IBLT of one set and the whole other set. To do this, we define *T*_*A*_ and *T*_*B*_ to be of the same size and to use the same hash functions. We then compute the difference of *T*_*A*_ and *T*_*B*_, denoted *T*_*A−B*_ and defined through *T*_*A−B*_[*i*].*C* = *T*_*A*_[*i*].*C* − *T*_*B*_[*i*].*C, T*_*A−B*_[*i*].*P* = *T*_*A*_[*i*].*P* ⊕ *T*_*B*_[*i*].*P*, and *T*_*A−B*_[*i*].*H* = *T*_*A*_[*i*].*H* ⊕ *T*_*B*_[*i*].*H*. Information about elements of *A* ∩ *B* is “cancelled out” in *T*_*A−B*_, that is, *T*_*A−B*_ holds elements of (*A* \ *B*) ∪ (*B* \ *A*). Peeling then proceeds as usual, listing both *A* \ *B* and *B* \ *A* with the distinction made possible by looking at the sign of *C*.

A remarkable property of this scheme is that it allows one to recover set differences using a space proportional to the size of those differences regardless the size of the involved sets. Indeed, for the peeling process to succeed w.h.p., it is sufficient that the size of *T*_*A−B*_ be *O*(*n*) where *n* = |(*A* \ *B*) ∪ (*B* \ *A*)| (see (1)). This is particularly suitable for the bioinformatics framework where we are often dealing with highly similar datasets, such as genomes of different individuals or closely related species.

### 3.2 Making buckets lighter

In the above scheme of IBLT difference, the *H* field becomes important as the case *T*_*A−B*_[*i*].*C* = 1 (or *T*_*A−B*_[*i*].*C* = −1) can occur due to a spurious “cancelling out” of distinct keys. However, to save space, we propose to get rid of the *H* field and replace the “checksum” verification by another test: if *T*_*A−B*_[*i*].*C* = 1 (resp. *T*_*A−B*_[*i*].*C* = −1), we check whether *p* = *T*_*A−B*_[*i*].*P* is a valid key by checking if *h*_*j*_(*p*) = *i* for one of *j* ∈ [1..*r*]. This allows us to save space at the price of additional verification time. This technique works particularly well for large IBLTs but it becomes less effective for small ones, as the “false positive” probability is proportional to the size of the table.

### 3.3 Combining sampling and IBLTs for Jaccard similarity estimation

We now turn to our main goal: estimating Jaccard similarity of two *k*-mer sets using IBLTs. The common approach uses MinHash sketching as described in [24] (see Section 2.4). However, MinHash requires larger sketches to measure similarity of close datasets. One possible idea could be to store MinHash sketches in IBLTs in hope to use them for estimating Jaccard similarity through the IBLT-difference scheme from the previous section. This, however, runs into an obstacle due to the fact that applying (2) requires knowledge of *k*-mers belonging to the sketch intersection, and not only to sketch differences.

Rather than working with the entire sets of *k*-mers, we resort to sampling. It is known that sampling minimizers incurs a bias in estimating Jaccard similarity [1]. Instead, we propose to use syncmers, which don’t suffer from being context-dependent thus resulting in an unbiased estimator of Jaccard similarity.

To justify the use of syncmers, we test a standard hash-based sampling, also providing an unbiased estimate of Jaccard similarity as well. To sample with a given sampling rate 1*/ν*, hash-based sampling uses a random hash function *h* : Σ^*k*^ → [0..*ν* − 1] with good statistical properties, and samples a *k*-mer *q* iff *h*(*q*) = 0.

Our approach consists in storing sampled *k*-mers in IBLTs and apply the IBLT-difference technique to recover set differences. Then, Jaccard similarity is estimated by

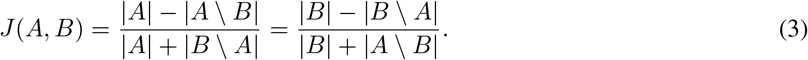

Note that cardinalities |*A*| and |*B*| can be easily retrieved from respective IBLTs *T*_*A*_ and *T*_*B*_ by summing all counter values and dividing by *r*.

### 3.4 IBLT dimensioning with syncmers

Dimensioning an IBLT holding syncmers requires estimating the expected number of differences in the set difference of involved *k*-mer sets. Assuming that input datasets are close genomic sequences of size *L* related by a mutation rate bounded by *p*_*m*_ and that *k* is sufficiently large so that *k*-mer occurrences are unique, we can estimate the set difference. Each mutation results in 2*k k*-mers in the set difference (*k k*-mers on each side), and therefore the size of set difference is estimated to be 2*kp*_*m*_*L*. Taking into account density 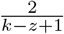 of syncmers (Section 2.2), we obtain the estimation

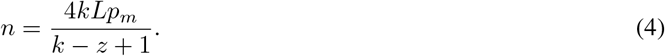

### 3.5 Approximating *k*-mer set differences

The method of Section 3.3 allows estimating Jaccard similarity on *k*-mers by Jaccard similarity on syncmers. Here we describe how we can extend these ideas in order to recover *all k*-mers from *K*(*S*_1_) \ *K*(*S*_2_) and *K*(*S*_2_) \ *K*(*S*_1_), where *S*_1_, *S*_2_ are input datasets and *K*(*S*) denotes the set of *k*-mers of a dataset *S*.

Note first that a straightforward way of doing this, through IBLTs of *K*(*S*_1_) and *K*(*S*_2_), requires a considerable space because a single mutation generates a difference of *k k*-mers. Using syncmers, we can “pack” *k*-mers into longer strings, compute the differences and then recover *k*-mers from them. The set of recovered *k*-mers, however, will be a superset of exact differences.

To achieve this, instead of storing syncmers, we store in IBLTs *extended* syncmers of length 2*k*−*z*. Extended syncmers are obtained by extending each syncmer to the right by *k* − *z* bases. Since successive syncmers overlap by at least *z* bases, this ensures that each *k*-mer belongs to at least one extended syncmer.

By applying the IBLT-difference technique (Section 3.3), we obtain the extended syncmers that differ between the two datasets, from which we extract *k*-mers and discard those shared between the two obtained sets. It may still happen that the sets we obtain are *supersets* of exact differences, due to the fact that an extended syncmer can contain a *k*-mer which belongs to another extended syncmer common to both datasets. However, we state that for a sufficiently large *k*, the fraction of common *k*-mers in those sets will be small enough, which we illustrate experimentally in Section 4.5. In the extreme case where each *k*-mer occurs once, our method computes exact *k*-mer set differences.

## 4 Results

To validate our ideas, we performed experiments on simulated sequences as well as on two real-life datasets:

- `covid`: subsample of 50 *SARS-CoV-2* genomes^1^. Sequence names are provided in Table 2.
- `spneu`: subsample of 28 *Streptococcus Pneumoniae* genomes from [5] whose names are reported in Table 3. The subsample has been chosen to contain very close strains, with pairwise mutation rates between them not exceeding 0.0005.

### 4.1 Comparison of different sampling approaches

Random sampling, minimizers and syncmers have been compared by computing Jaccard similarities between pairs of synthetic sequences. Each pair is constructed by first generating a uniform random sequence of length *L* and then mutating it through independent substitutions. Points in Figure 1a are averages over *T* = 500 independent trials. For fairness of comparison, parameters for uniform sampling, minimizers and syncmers have been chosen to guarantee the same sampling rate 1*/ν*. We know that *c*_*s*_ ≈ 2*/*(*k* − *z* + 1), *c*_*m*_ ≈ 2*/*(*w* + 1) and *c*_*ns*_ ≈ 1*/ν* are the densities of syncmers, minimizers and random sampling, respectively. Thus, given parameters *k* and *z*, setting the minimizer window length as *w* = *k*−*z* and choosing a sampling rate 1*/ν* = *c*_*s*_ ensures about the same number of sampled *k*-mers for all algorithms. As Figure 1a shows, syncmers do not have the previously reported biased behaviour of minimizers [1], but they seem to be comparable to random sampling. However, as shown in Figure 1b, random sampling is subject to larger errors than syncmers, due to less uniform distribution along the sequence. For these reasons, we choose syncmer sampling as the mean to reduce IBLT memory in Section 4.3.

**Figure 1:**
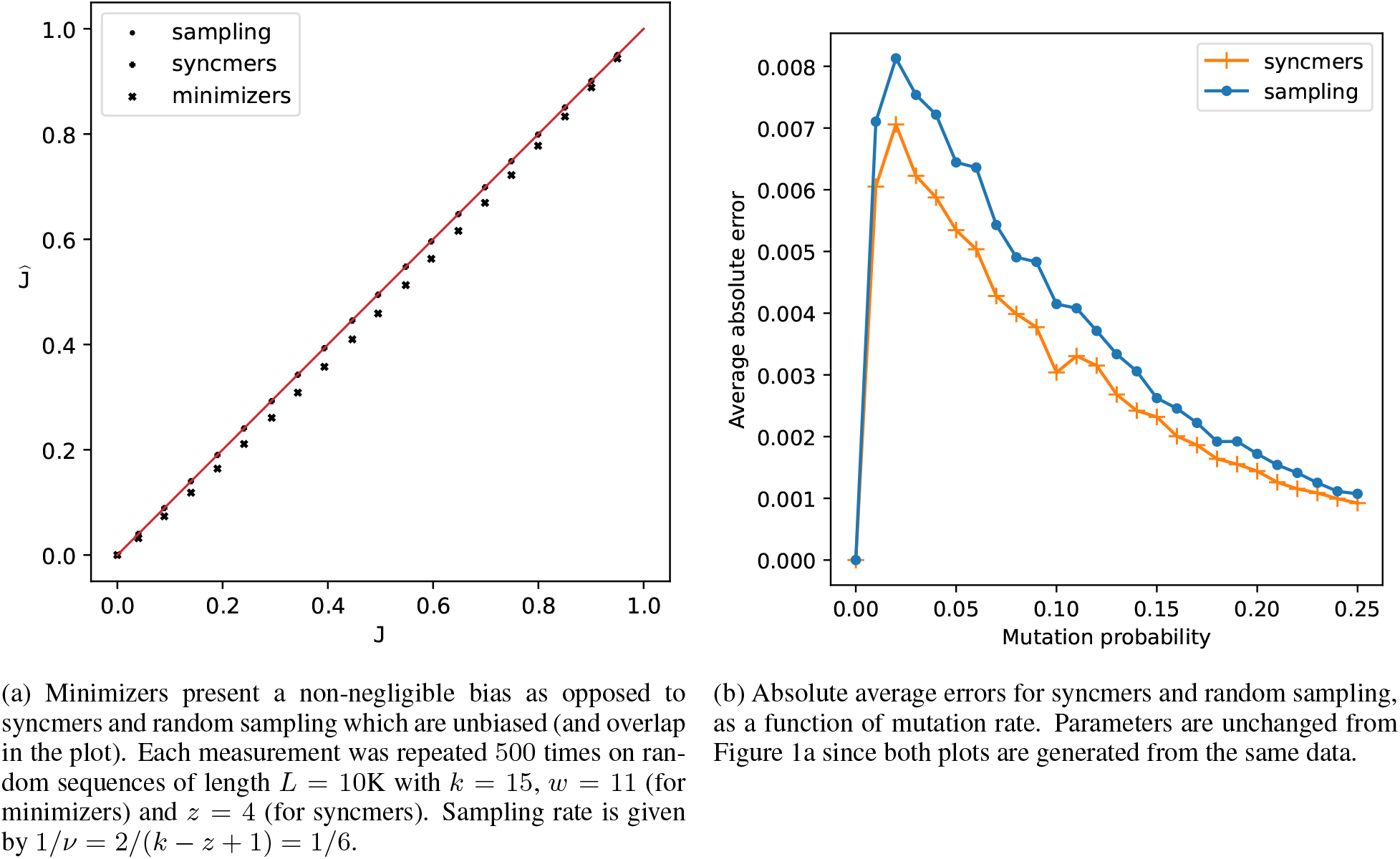
Comparison between random sampling, minimizers and syncmers.

### 4.2 Space performance of IBLTs

In order to demonstrate the space efficiency of IBLTs in our framework, we compare them against a solution based on KMC *k*-mer counting software [17]. KMC provides an efficient way for storing, manipulating and querying sets of *k*-mers. Unlike other counting tools (Jellyfish [21] or DSK [27]), KMC allows easy sorting of its output which leads to an efficient way to compute Jaccard similarity.

We compared memory taken by IBLTs vs. KMC databases for storing syncmers issued from two similar sequences. For this, we applied the same procedure as in Section 4.1: mutating a random sequence of length *L* with mutation probability *p*_*m*_. Sampled syncmers from both sequences are stored respectively in IBLTs and KMC databases. Figure 2 reports average space taken by the two data structures. Each bar is the average over *T* = 100 trials, except for case *L* = 10M for which *T* = 10. IBLTs were dimensioned (see (1)) to guarantee peelability of all *T* sketches with high probability.

**Figure 2:**
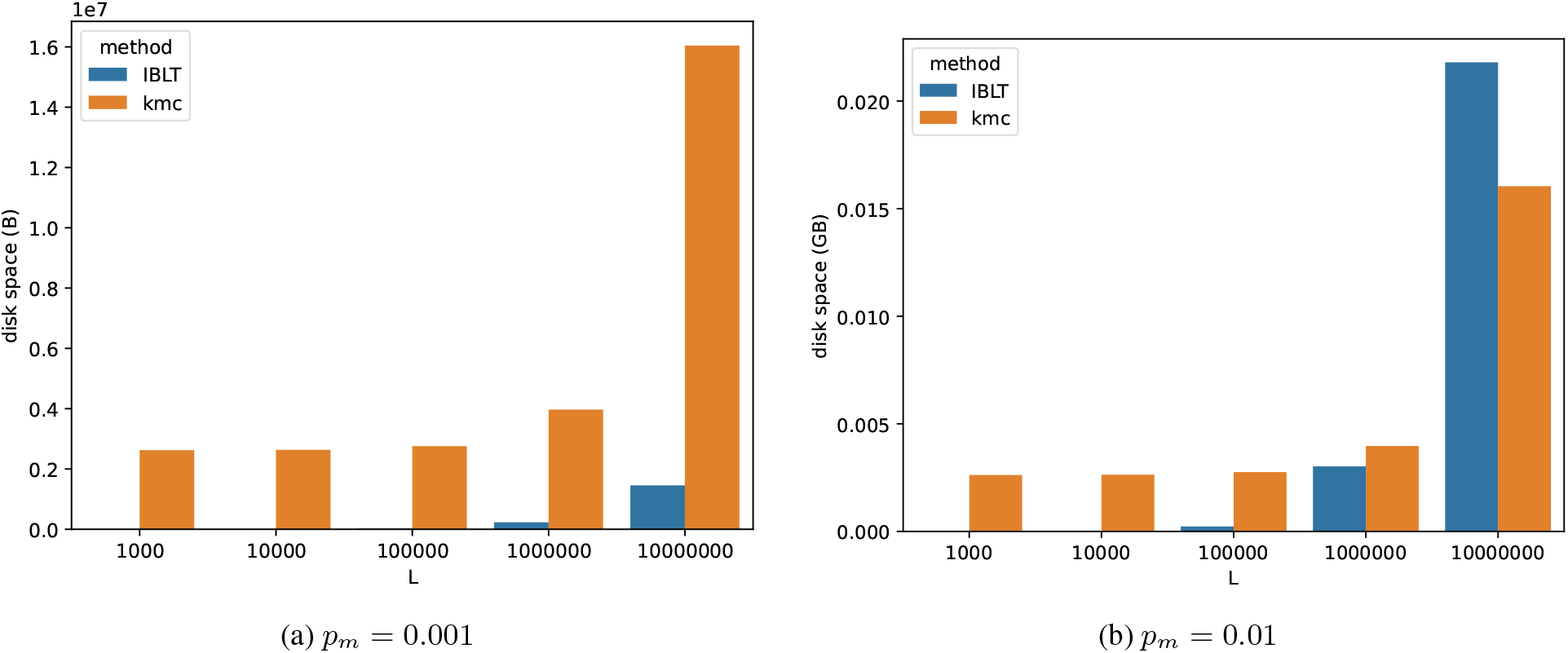
Space taken by IBLTs depends on the similarity between stored sets. For very similar sequences (mutation rate *p*_*m*_ = 0.001, Figure 2a), IBLTs are more space-efficient than KMC. Their advantage appears reduced for increased *p*_*m*_ and large sequences (Figure 2b).

Figure 2a clearly demonstrates the advantage of IBLTs when the mutation rate is small. For larger *p*_*m*_ and long sequences, the number of differences reach a point where exact data structures become preferable, as illustrated by Figure 2b for *p*_*m*_ = 0.01 and sequences of length 10M.

In our experiments, subtracting one IBLTs from another is dominated by the time taken to load/save the sketches, and not by performing the actual difference. Even in more complex scenarios, subtraction remains a very simple operation that can be performed by accessing one bucket at a time in any given order. On the other hand, the amount of time required by listing the content of an IBLT varies greatly and depends on the set of items stored in it.

### 4.3 Accuracy of Jaccard similarity estimation from IBLTs of syncmers

Figures 3 and 4 report comparisons of both IBLTs and MinHash sketches on `covid` and `spneu` datasets respectively. Both plots show the average absolute error of Jaccard estimate computed over all pairs of sequences of the respective dataset. Exact Jaccard similarities computed over the full *k*-mer sets are used as ground truth. MinHash sketches (line MINHASH in the plots) were implemented using MASH [24]. All sketch sizes (in bytes) are fixed beforehand with both MinHash sketches and IBLTs dimensioned accordingly in order to fit the allocated memory. The number of bits allocated for payload field *P* in our IBLT implementation is set to be the minimum multiple of 8 larger than or equal to 2*k*. As MASH [24] uses 32-or 64-bit hashes, we used *k* = 15 in our experiments in order to force both methods to use 32-bit representations.

**Figure 3:**
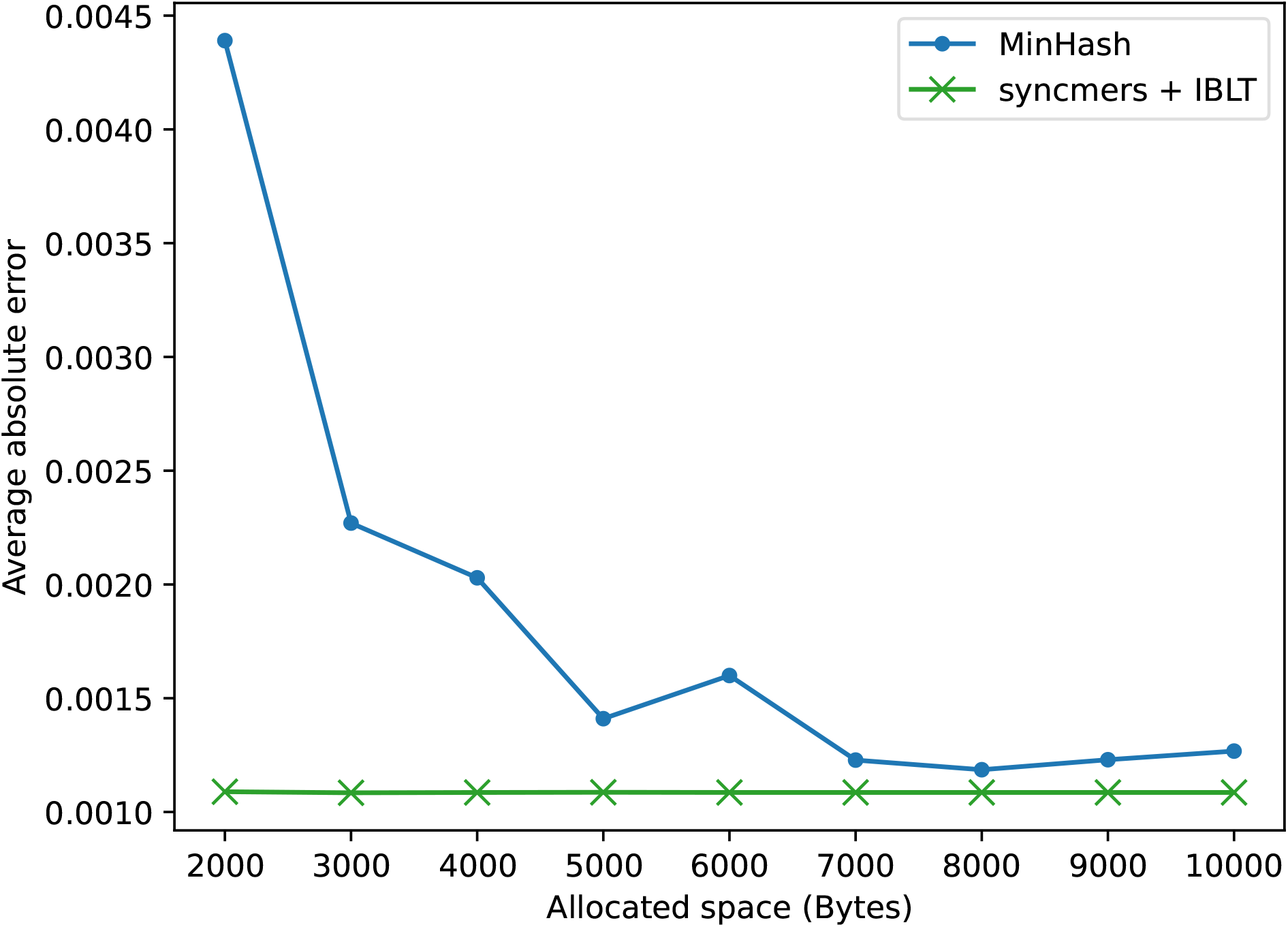
Comparison between IBLTs and MinHash for computing pairwise Jaccard on the `covid` dataset. The x-axis reports the amount of space allocated for each sketch while the y-axis reports the average absolute error. *k* = 15 and *z* = 4 in all tests. Sketch size for MinHash and table size for IBLTs are chosen to fit the allocated memory.

**Figure 4:**
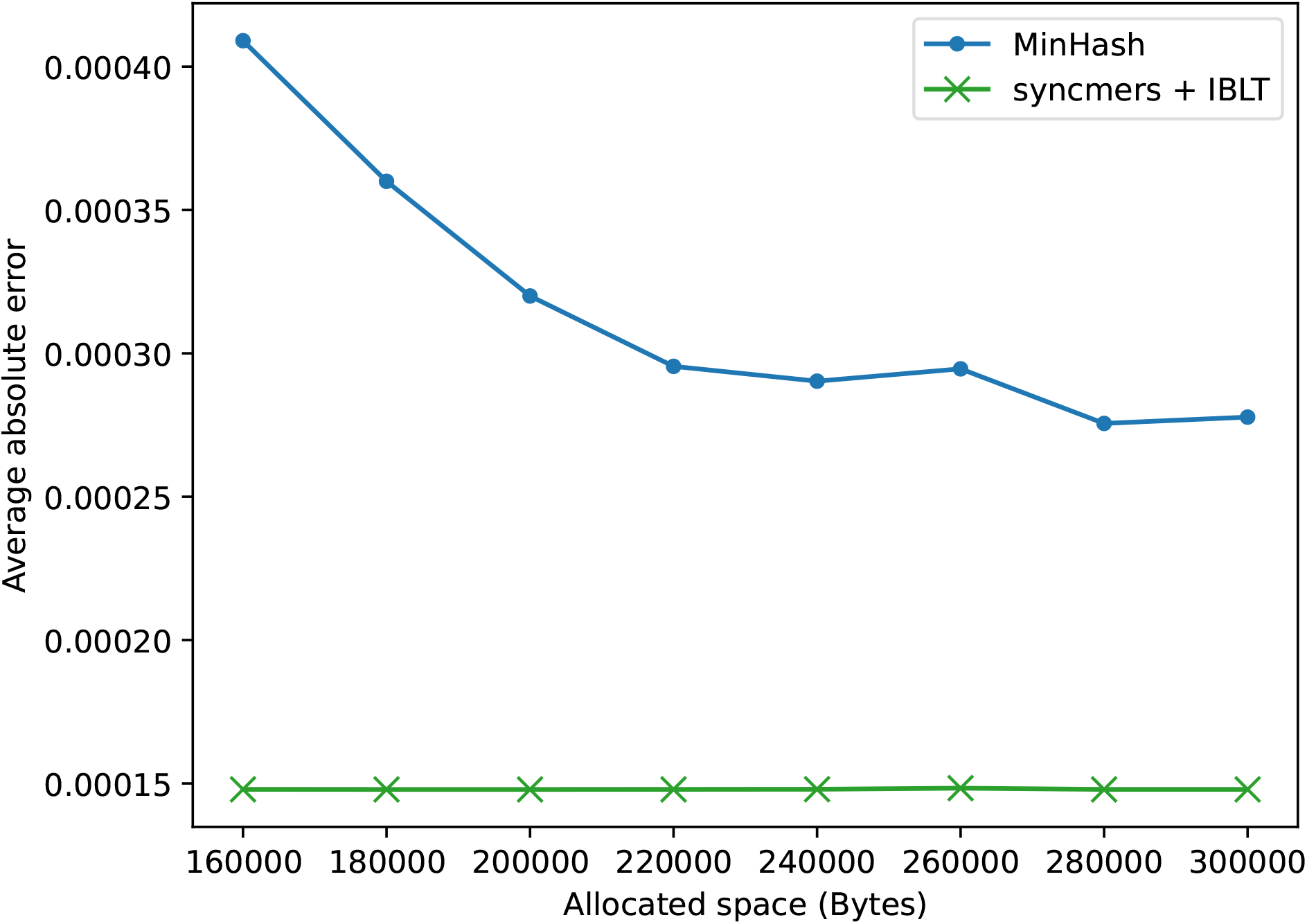
Comparison between IBLTs and MinHash for computing pairwise Jaccard on the `spneu` dataset with the same setting as Figure 3.

In all experiments, IBLTs storing syncmers (line SYNCMERS + IBLT) showed the best precision. For covid genomes (Figure 3), full MinHash sketches become competitive for larger sketch sizes. Unlike MinHash, the average error of IBLTs remains constant across all reported cases because over-dimensioning only increases the probability of successful listing. For the `spneu` dataset (Figure 4), MinHash errors are about twice those of IBLTs across all allocated sketch sizes confirming that IBLTs are more memory-efficient. The general conclusion is that if sequences to be compared are highly similar, IBLTs storing syncmers are more efficient than MinHash sketches, with the latter being better suited to quickly provide an overview over more diverging datasets.

### 4.4 Sampling syncmers for further space reductions

Since syncmer sampling rate 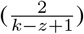 cannot be made arbitrarily small for a given *k*, we also tested the effect of additional downstream sampling of syncmers, before inserting them into IBLTs. To this end, Figure 5 reports a comparison of syncmers sampled with different sampling rates 1*/ν*. We observe that downstream sampling of syncmers comes at the cost of decreased precision for both datasets (Figure 5a and 5b), but it might be useful to further reduce space.

**Figure 5:**
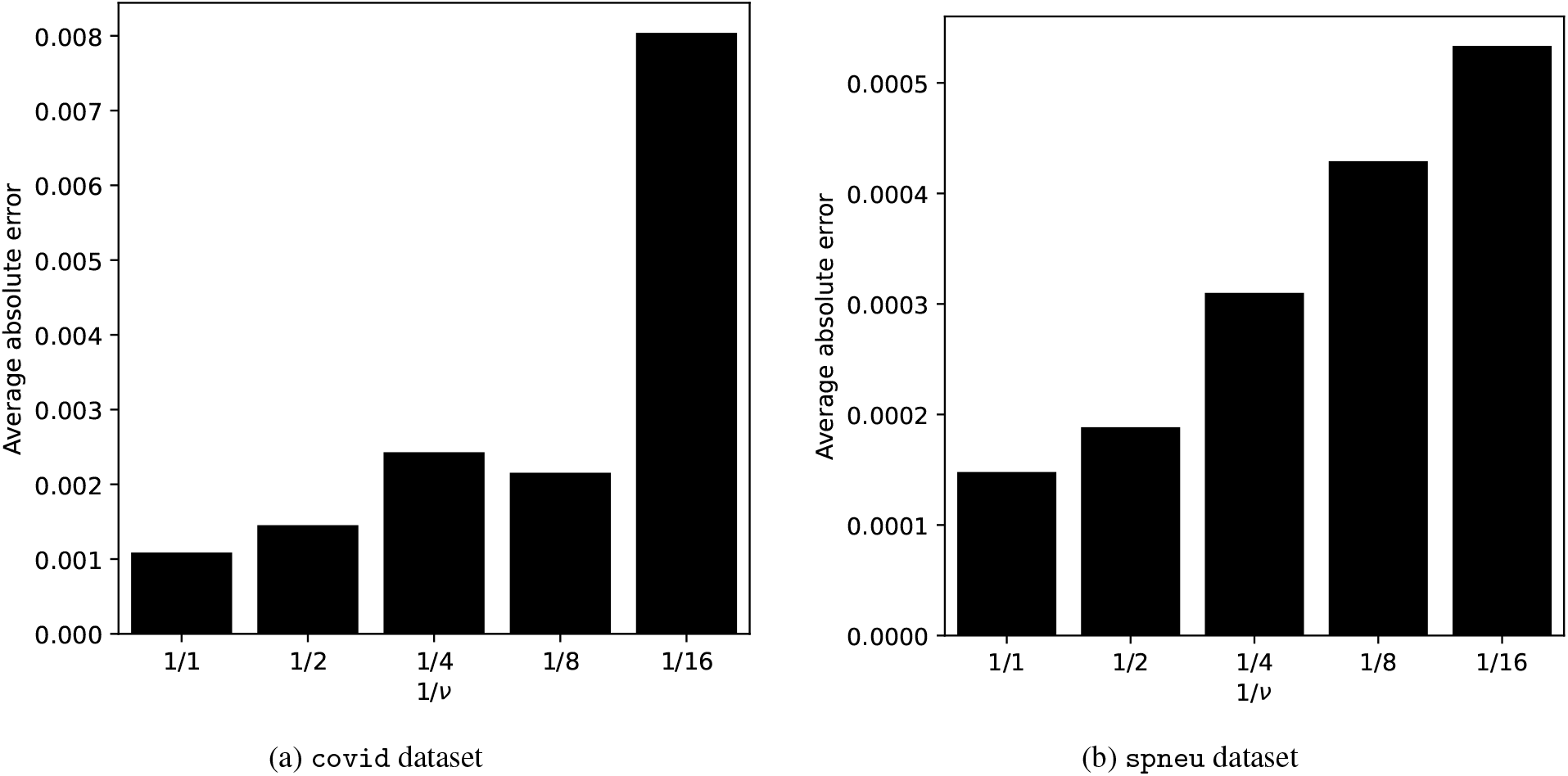
Effect of sampling syncmers before IBLT insertion on the average absolute error. 1*/ν* is the compression rate used for sampling syncmer sets before IBLT insertion. *ν* = 1 means no sampling (full syncmer sets).

### 4.5 Experiments on approximating *k*-mer set differences

We tested the method from Section 3.5 of approximating *k*-mer set differences on both the `covid` dataset and on two random datasets. Each random dataset contains 50 sequences of length 30000 obtained by first generating a uniform random sequence which is then mutated 49 times using a mutation probability *p*_*m*_.

Recall that the method of Section 3.5 allows one to compute a *superset* of the symmetric difference (*K*(*S*_1_)\*K*(*S*_2_))∪ (*K*(*S*_2_) \ *K*(*S*_1_)) of sets of *k*-mers occurring in datasets *S*_1_ and *S*_2_. Here we measure the precision of this method, that is the number of spurious *k*-mers found by the algorithm. Those are *k*-mers actually belonging to *K*(*S*_1_) ∩ *K*(*S*_2_) but output by the algorithm as if they belong to (*K*(*S*_1_) \ *K*(*S*_2_)) ∪ (*K*(*S*_2_) \ *K*(*S*_1_)).

Table 1 summarizes the experiments. Columns ‘diff’ and ‘err’ show the average/maximum cardinality of the true set difference and spurious *k*-mers, respectively, over all pairs of sequences. In the case of random datasets, sequences were generated with mutation probabilities *p*_*m*_ = 0.01 and *p*_*m*_ = 0.001.

**Table 1:**
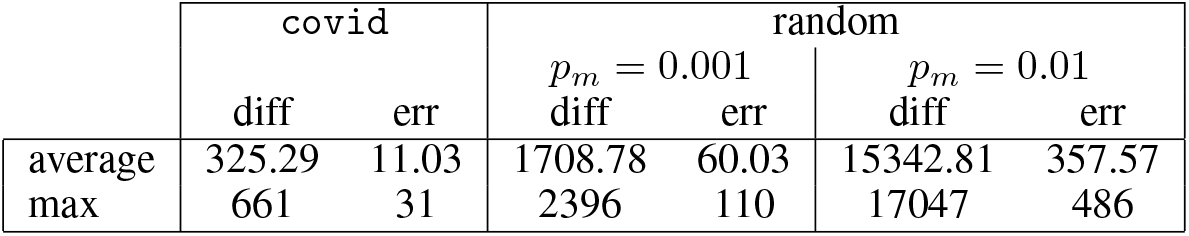
True size of symmetric difference of *k*-mer sets and its overestimate. For each experiment, ‘diff’ is the average/maximum size of the true symmetric difference, and ‘err’ is the average/maximum number of spurious *k*-mers reported as being in the symmetric difference. *p*_*m*_ is the mutation probability used to generate sequences from a random one.

|  | covid |  | random |  |  |  |
| --- | --- | --- | --- | --- | --- | --- |
| | diff | err | $p_m = 0.001$<br>diff | $p_m = 0.001$<br>err | $p_m = 0.01$<br>diff | $p_m = 0.01$<br>err |
| average | 325.29 | 11.03 | 1708.78 | 60.03 | 15342.81 | 357.57 |
| max | 661 | 31 | 2396 | 110 | 17047 | 486 |

We observe that the number of spurious *k*-mers remains small, on average within about 3% of the true set difference size.

## 5 Conclusions

To the best of our knowledge, our work is the first to apply Invertible Bloom Lookup Tables to *k*-mer processing for alignment-free comparison of DNA sequence datasets. We showed that whenever involved datasets are similar enough and their similarity can be bounded *a priori*, IBLTs lead to a more space-efficient and, at the same time, more accurate method for estimating Jaccard similarity of underlying *k*-mer sets. This is achieved by combining IBLTs with *k*-mer sampling via syncmers. As opposed to minimizers, syncmers provide an unbiased estimator of Jaccard index, which was confirmed in our experiments. At the same time, syncmer sampling is shown to lead to a more concentrated estimator than the straightforward hash-based sampling. Thus, IBLTs combined with syncmers constitute a powerful alternative to MinHashing for estimating Jaccard similarity for similar datasets. Note that in the context of viral/bacterial pan-genomics, dealing with similar datasets is a predominant situation in bioinformatics. In particular, the number of closely related bacterial and viral strains is rapidly growing.

As another application of IBLTs, we are able to approximately compute differences of underlying *k*-mer sets in small space. This opens new prospects as *k*-mers proper to a dataset can be used to infer information about genetic variation, specific mutation, etc. Note that MINHASH is designed to only estimate similarity and is not capable of providing information about actual differences. We also believe that by using additional space-efficient data structures this method can be extended to compute *exact* set differences on more complex datasets and we plan to explore this in our future work.

Our ideas may have further useful applications, for example to reconciliation of datasets located on remote computers, in which case IBLTs could avoid transmitting entire datasets (similar to a scenario described in [12]). Another example is a selection of sufficiently diverse datasets avoiding redundancy, as e.g. [18]. Note finally that IBLTs may also act as filters for filtering out dissimilar datasets: in this case, non-peelability of the difference IBLT is an indicator of dissimilarity.

## 6 Appendix

### Datasets

**Table 2:** Names of covid genomes used for Figure 3

|  |  |  |  |
| --- | --- | --- | --- |
| BS001151.1 | LR877722.1 | LR883214.1 | MT520216.1 |
| MT706180.1 | MT757082.1 | MT800758.1 | MT834020.1 |
| MT970159.1 | MT971010.1 | MT973151.1 | MW064390.1 |
| MW064919.1 | MW064981.1 | MW153809.1 | MW153954.1 |
| MW154711.1 | MW156712.1 | MW184416.1 | MW184648.1 |
| MW190904.1 | MW190957.1 | MW191020.1 | MW191146.1 |
| MW206148.1 | MW276931.1 | MW321243.1 | MW321430.1 |
| MW593629.1 | MW631874.1 | MW669599.1 | MW681303.1 |
| MW681489.1 | MW693959.1 | MW696216.1 | MW702101.1 |
| MW708072.1 | MW708184.1 | MW708826.1 | MW720341.1 |
| MW733722.1 | MW738615.1 | MW749542.1 | MW776764.1 |
| MW820211.1 | MW850083.1 | MW863243.1 | MW868532.1 |
| MW868533.1 | MW871079.1 |  |  |

**Table 3:**
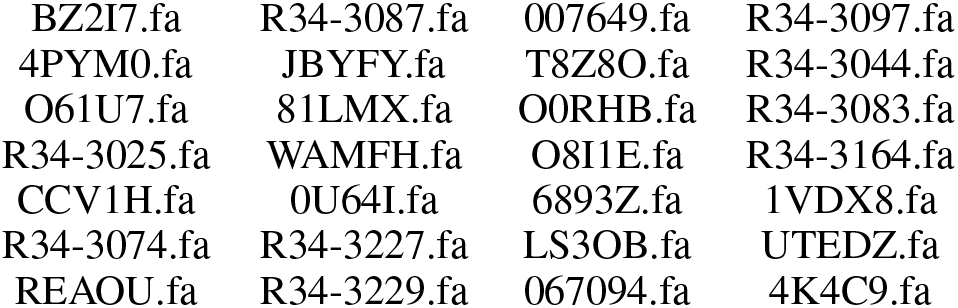
Names of *S*.*Pneumoniae* genomes used for Figure 4

## Footnotes

1 https://www.ncbi.nlm.nih.gov/datasets/coronavirus/genomes

## References

[1] Mahdi Belbasi, Antonio Blanca, Robert S. Harris, David Koslicki, and Paul Medvedev. The minimizer Jaccard estimator is biased and inconsistent. bioRxiv, 2022. arXiv:https://www.biorxiv.org/content/early/2022/01/17/2022.01.14.476226.full.pdf, doi:10.1101/2022.01.14.476226.

[2] Timo Bingmann, Phelim Bradley, Florian Gauger, and Zamin Iqbal. COBS: A Compact Bit-Sliced Signature Index. In Nieves R. Brisaboa and Simon J. Puglisi, editors, String Processing and Information Retrieval, Lecture Notes in Computer Science, pages 285–303, Cham, 2019. Springer International Publishing. doi:10.1007/978-3-030-32686-9_21.

[3] Phelim Bradley, Henk C Den Bakker, Eduardo P. C. Rocha, Gil McVean, and Zamin Iqbal. Ultra-fast search of all deposited bacterial and viral genomic data. Nature biotechnology, 37(2):152–159, February 2019. URL: https://www.ncbi.nlm.nih.gov/pmc/articles/PMC6420049/, doi:10.1038/s41587-018-0010-1.

[4] Andrei Z. Broder. On the resemblance and containment of documents. In Proceedings. Compression and Complexity of SEQUENCES 1997 (Cat. No.97TB100171), pages 21–29, June 1997. doi:10.1109/SEQUEN.1997.666900.

[5] Karel Břinda, Alanna Callendrello, Kevin C. Ma, Derek R. MacFadden, Themoula Charalampous, Robyn S. Lee, Lauren Cowley, Crista B. Wadsworth, Yonatan H. Grad, Gregory Kucherov, Justin O’Grady, Michael Baym, and William P. Hanage. Rapid inference of antibiotic resistance and susceptibility by genomic neighbour typing. Nature Microbiology, 5(3):455–464, March 2020. URL: https://www.nature.com/articles/s41564-019-0656-6, doi:10.1038/s41564-019-0656-6.

[6] Dan DeBlasio, Fiyinfoluwa Gbosibo, Carl Kingsford, and Guillaume Marçais. Practical universal k-mer sets for minimizer schemes. In Proceedings of the 10th ACM International Conference on Bioinformatics, Computational Biology and Health Informatics, BCB ‘19, page 167–176, New York, NY, USA, 2019. Association for Computing Machinery. doi:10.1145/3307339.3342144.

[7] Martin Dietzfelbinger, Andreas Goerdt, Michael Mitzenmacher, Andrea Montanari, Rasmus Pagh, and Michael Rink. Tight thresholds for cuckoo hashing via xorsat. In Proceedings of the 37th International Colloquium Conference on Automata, Languages and Programming, ICALP’10, page 213–225, Berlin, Heidelberg, 2010. Springer-Verlag.

[8] Robert Edgar. Syncmers are more sensitive than minimizers for selecting conserved k-mers in biological sequences. PeerJ, 9:e10805, February 2021. URL: https://peerj.com/articles/10805, doi:10.7717/peerj.10805.

[9] Barış Ekim, Bonnie Berger, and Yaron Orenstein. A Randomized Parallel Algorithm for Efficiently Finding Near-Optimal Universal Hitting Sets. In Russell Schwartz, editor, Research in Computational Molecular Biology, Lecture Notes in Computer Science, pages 37–53, Cham, 2020. Springer International Publishing. doi:10.1007/978-3-030-45257-5_3.

[10] David Eppstein and Michael T. Goodrich. Straggler identification in round-trip data streams via Newton’s identities and invertible Bloom filters. IEEE Transactions on Knowledge and Data Engineering, 23(2):297–306, 2011. doi:10.1109/TKDE.2010.132.

[11] Huan Fan, Anthony R. Ives, Yann Surget-Groba, and Charles H. Cannon. An assembly and alignment-free method of phylogeny reconstruction from next-generation sequencing data. BMC Genomics, 16(1):522, July 2015. doi:10.1186/s12864-015-1647-5.

[12] Michael T. Goodrich and Michael Mitzenmacher. Invertible Bloom lookup tables, 2011. URL: https://arxiv.org/abs/1101.2245, doi:10.48550/ARXIV.1101.2245.

[13] Gaurav Gupta, Minghao Yan, Benjamin Coleman, R. A. Leo Elworth, Todd Treangen, and Anshumali Shrivastava. Sub-linear Sequence Search via a Repeated And Merged Bloom Filter (RAMBO): Indexing 170 TB data in 14 hours. arXiv:1910.04358 [cs, q-bio], December 2019. arXiv: 1910.04358. URL: http://arxiv.org/abs/1910.04358.

[14] Robert S Harris and Paul Medvedev. Improved representation of sequence Bloom trees. Bioinformatics, 36(3):721–727, 08 2019. arXiv:https://academic.oup.com/bioinformatics/article-pdf/36/3/721/38712473/btz662.pdf, doi:10.1093/bioinformatics/btz662.

[15] Chirag Jain, Luis M. Rodriguez-R, Adam M. Phillippy, Konstantinos T. Konstantinidis, and Srinivas Aluru. High throughput ANI analysis of 90K prokaryotic genomes reveals clear species boundaries. Nature Communications, 9(1):5114, November 2018. URL: https://www.nature.com/articles/s41467-018-07641-9, doi:10.1038/s41467-018-07641-9.

[16] Mikhail Karasikov, Harun Mustafa, Gunnar Rätsch, and André Kahles. Lossless indexing with counting de bruijn graphs. bioRxiv, 2022. arXiv:https://www.biorxiv.org/content/early/2022/02/03/2021.11.09.467907.full.pdf, doi:10.1101/2021.11.09.467907.

[17] Marek Kokot, Maciej Długosz, and Sebastian Deorowicz. KMC 3: counting and manipulating k-mer statistics. Bioinformatics, 33(17):2759–2761, 05 2017. arXiv:https://academic.oup.com/bioinformatics/article-pdf/33/17/2759/25163903/btx304.pdf, doi:10.1093/bioinformatics/btx304.

[18] Nathan LaPierre, Mohammed Alser, Eleazar Eskin, David Koslicki, and Serghei Mangul. Metalign: efficient alignment-based metagenomic profiling via containment min hash. Genome Biology, 21(1):242, September 2020. doi:10.1186/s13059-020-02159-0.

[19] Heng Li. Minimap2: pairwise alignment for nucleotide sequences. Bioinformatics, 34(18):3094–3100, 05 2018. arXiv:https://academic.oup.com/bioinformatics/article-pdf/34/18/3094/25731859/bty191.pdf, doi:10.1093/bioinformatics/bty191.

[20] Camille Marchet, Christina Boucher, Simon J. Puglisi, Paul Medvedev, Mikaël Salson, and Rayan Chikhi. Data structures based on k-mers for querying large collections of sequencing data sets. Genome Research, 31(1):1–12, January 2021. URL: https://www.ncbi.nlm.nih.gov/pmc/articles/PMC7849385/, doi:10.1101/gr.260604.119.

[21] Guillaume Marçais and Carl Kingsford. A fast, lock-free approach for efficient parallel counting of occurrences of k-mers. Bioinformatics, 27(6):764, March 2011. doi:10.1093/bioinformatics/btr011.

[22] Michael Molloy. The pure literal rule threshold and cores in random hypergraphs. In Proceedings of the fifteenth annual ACM-SIAM symposium on Discrete algorithms, SODA ‘04, pages 672–681, USA, January 2004. Society for Industrial and Applied Mathematics.

[23] Martin D Muggli, Bahar Alipanahi, and Christina Boucher. Building large updatable colored de Bruijn graphs via merging. Bioinformatics, 35(14):i51–i60, 07 2019. arXiv:https://academic.oup.com/bioinformatics/article-pdf/35/14/i51/28913259/btz350.pdf, doi:10.1093/bioinformatics/btz350.

[24] Brian D. Ondov, Todd J. Treangen, Páll Melsted, Adam B. Mallonee, Nicholas H. Bergman, Sergey Koren, and Adam M. Phillippy. Mash: fast genome and metagenome distance estimation using MinHash. Genome Biology, 17(1):132, June 2016. doi:10.1186/s13059-016-0997-x.

[25] Giulio Ermanno Pibiri. Sparse and skew hashing of k-mers. bioRxiv, 2022. arXiv:https://www.biorxiv.org/content/early/2022/01/18/2022.01.15.476199.full.pdf, doi:10.1101/2022.01.15.476199.

[26] Ely Porat and Ohad Lipsky. Improved sketching of hamming distance with error correcting. In Bin Ma and Kaizhong Zhang, editors, Combinatorial Pattern Matching, pages 173–182, Berlin, Heidelberg, 2007. Springer Berlin Heidelberg.

[27] Guillaume Rizk, Dominique Lavenier, and Rayan Chikhi. DSK: k-mer counting with very low memory usage, March 2013. doi:10.1093/bioinformatics/btt020.

[28] Michael Roberts, Wayne Hayes, Brian R. Hunt, Stephen M. Mount, and James A. Yorke. Reducing storage requirements for biological sequence comparison. Bioinformatics, 20(18):3363–3369, December 2004. doi: 10.1093/bioinformatics/bth408.

[29] Kamil Salikhov, Gustavo Sacomoto, and Gregory Kucherov. Using cascading Bloom filters to improve the memory usage for de Brujin graphs. BMC Algorithms for Molecular Biology, 9(1):2, 2014. URL: http://www.almob.org/content/9/1/2.

[30] Saul Schleimer, Daniel S. Wilkerson, and Alex Aiken. Winnowing: local algorithms for document fingerprinting. In Proceedings of the 2003 ACM SIGMOD international conference on Management of data, SIGMOD ‘03, pages 76–85, San Diego, California, June 2003. Association for Computing Machinery. doi:10.1145/872757.872770.

